# Doxorubicin-Induced p53 Interferes with Mitophagy in Cardiac Fibroblasts

**DOI:** 10.1101/674309

**Authors:** TR Mancilla, GJ Aune

**Author notes:** Address for Correspondence: Gregory J Aune, MD, PhD, 8403 Floyd Curl Drive, San Antonio, TX 78229-3900.

## Abstract

Doxorubicin is a mainstay in pediatric chemotherapy treatment because of its efficacy treating leukemia and lymphoma. Unfortunately, every childhood cancer survivor will develop a chronic health problem, one of the most serious being cardiac disease. How doxorubicin damages the heart in such a way that disease progression occurs over multiple decades is still not understood.

The dose of doxorubicin selected does not cause apoptosis but does arrest cell cycle. It also decreases the cells ability to migrate. Gene profiling indicated a cardiac remodeling and inflammatory profile. Mitochondria increased ROS production and underwent membrane depolarization. Secondly, the Parkin:p53 interaction mechanism was investigated. Doxorubicin was found to increase p53 expression and it was shown to sequester Parkin. As a result, mitophagy in doxorubicin-treated cells was decreased. Lastly, cardiac fibroblasts were isolated from p53^-/-^ mice and treated with doxorubicin. The gene expression phenotype in these cells was attenuated and migration was restored. Proliferation was still decreased. Mitochondrial dysfunction was also partially attenuated. Without p53, Parkin could now localize to the mitochondria and mitophagy was restored.

Doxorubicin induces a deleterious phenotype in cardiac fibroblasts that may be due to the interaction between two stress responses caused by doxorubicin’s DNA and mitochondrial damage. Cardiac fibroblasts are a viable target and further research needs to be done to elucidate other harmful mechanisms at play in the fibroblast. Knowledge about the importance of cardiac fibroblasts in the development of doxorubicin-induced cardiotoxicity and a pathological mechanism broadens our understanding and ability to develop protective therapies to improve the quality of life of cancer survivors.

The project described was supported by all of the following sources for GJA:

- St. Baldrick’s Foundation Scholar (Career Development Award)
- Turn it Gold Foundation

The project described was supported by all of the following sources for TRM:

- NIH T32GM113896 (STX-MSTP) award
- National Center for Advancing Translational Science, NIH through grant TL1 TR001119. The content is solely the responsibility of the authors and does not necessarily represent the official views of the NIH.

## Introduction

By age 50, over half of all childhood cancer survivors will suffer from at least one debilitating or even fatal chronic health condition^1^. Moreover, by 2020, due to improved treatment regimens, there will be over 500,000 childhood cancer survivors in the U.S. alone^2^. The most common fatal late complications in these survivors are secondary neoplasms and heart disease^3^. Epidemiological data has unequivocally linked doxorubicin (DOX), a chemotherapy agent used in over 50% of pediatric cancer cases^4^, to acute and chronic cardiotoxicity^5–7^. While the drug has been in use since the 1970’s, clinical and preclinical research has yet to produce an effective strategy to prevent chronic cardiotoxicity before or after exposure.

A challenging barrier to understanding the pathologic mechanisms of chronic DOX cardiotoxicity is the latent period. In patients that are treated during childhood, cardiac dysfunction may not be detectable for two or three decades. DOX toxicity manifests differently in cancer survivors treated as adults from those treated as children. Childhood cancer survivors suffer from chronic cardiotoxicity, whereas acute toxicity has almost been eliminated in these patients due to cumulative dose limits^8^. Adult survivors have a much higher incidence of acute cardiotoxicity. Adult survivors are also prone to chronic cardiotoxicity, but it can be difficult to separate from the natural cardiac dysfunction of aging^9^. There are also molecular and physiological differences contributing to the divergence in pediatric and adult patient outcomes. At the time of their chemotherapy, children are still undergoing cardiac maturation, the process whereby the cardiac myocytes hypertrophy to match the increasing demands of the growing body^10^. Dysfunction to the various cells of the heart due to DOX exposure could lead to harmful alterations in the maturation process. Cardiac fibroblasts, in particular, use paracrine signaling to mediate cardiac myocyte hypertrophy^11–13^ and ECM remodeling^14; 15^ necessary for maturation.

The anti-tumor mechanism of action of DOX is imparted via DNA intercalation and inhibition of topoisomerase II. DOX then disrupts DNA and macromolecular synthesis and triggers double strand DNA breaks^16; 17^. This mechanism is thought to target tumor cells over other cells, especially non-proliferating and mitotically inactive cells. Physicians and researchers initially believed that what made DOX such an effective cancer agent was separate from cardiotoxicity. However, there may not be a clear delineation between the two processes. Prior research has established the severe pathological processes triggered by DOX in cardiac myocytes. The prevailing theory for myocyte damage centers on DOX induction of reactive oxygen species (ROS)^7^, which leads to mitochondrial damage and apoptosis^18^. Due to high mitochondrial load and a lack of scavenger molecules, cardiac myocytes are more prone to oxidative damage than other cell types^19^.

In addition to myocytes, cardiac fibroblasts are key players in myocardial stress responses, including cardiac extracellular remodeling and wound healing^20–22^. Cardiac fibroblasts interact with myocytes through direct connections and paracrine signaling^23–26^. In a maturing heart, cardiac fibroblasts also regulate cardiac myocyte growth^27; 28^. Dysfunction of cardiac fibroblasts in a pediatric patient could cause a progressive pathology and yet, little is known about the direct effects of DOX on cardiac fibroblasts. A recent study demonstrated the importance of cardiac fibroblasts in DOX-induced cardiac dysfunction via cardiac fibroblast-specific knockdown of the DNA damage response gene *Ataxia telangiectasia mutated* (ATM) kinase^29^. Cardiac fibroblast-specific changes were able to mitigate DOX-induced cardiotoxicity. Fibroblasts were studied as opposed to myocytes, because the ATM signal was predominantly in fibroblasts.

Just as in other cells, mitochondria are critical regulators of cardiac fibroblast health. Research into mitochondrial health after DOX exposure has not been conducted in the cardiac fibroblast. We do know, from research on the cardiac myocyte that DOX can induce mitochondrial dysfunction and depolarization. Mitophagy, the degradation and recycling of mitochondria, is one way to maintain mitochondrial health. A recent study has shown that p53, which is upregulated in cardiac cells after exposure to DOX, can inhibit mitophagy by sequestering Parkin in the cytosol^30^. Normally, Parkin locates to damaged mitochondria to initiate autophagosome engulfment, an early step of mitophagy. Without the ability to undergo mitophagy, a cell accumulates dysfunctional mitochondria perpetuating ROS production^31; 32^. What is unknown is how cardiac fibroblasts respond to DOX-induced stress, such as mitochondrial dysfunction. Over time, these damaged mitochondria could create a cardiac milieu primed for adverse remodeling that would then gradually progress to overt heart failure.

A 2005 study by *Shizukuda et al*. compared cardiac function after DOX exposure in WT C57BL/6J mice and p53 KO mice^33^. While the model used only a single large injection, the KO mice showed no change in cardiac function for up to two weeks after treatment. Cardiac dysfunction via a decrease in ejection fraction was evident within four days of treatment in the WT mice and a statistically and physiologically significant decrease was seen at two weeks follow up. *Hoshido et al*. determined that cytosolic p53 directly binds to Parkin preventing its transportation to the mitochondria^30^. This was demonstrated *in vitro* using mesenchymal embryonic fibroblasts (MEFs) and the rescue of mitophagy in p53^-/-^ MEFs. The association and rescue of mitophagy was also shown in WT and p53^-/-^ mice treated with DOX. Two findings indicated that the Parkin:p53 interaction was upstream of mitophagy. In p53^-/-^ mice treated with DOX, there was an increase in the number of mitochondria engulfed by autophagosomes. Double knockout mice of Parkin and p53 could not rescue the mitophagy process after DOX treatment.

Interestingly, *Feridooni et al*. repeated the work of Shizukuda, but used a p53^-/-^ restricted to cardiac myocytes. In this model the previous benefits of p53 knockdown (KD) were no longer seen^34^. The authors concluded that the benefits seen with p53 KD previously were p53-independent. They did not question whether the role of p53 could be significant in another cell type.

Oxidative stress in fibroblasts has been investigated in other tissue types, especially skin fibroblasts. One study showed that exogenous or increased endogenous ROS could upregulate MMP expression and downregulate collagen expression^35^. Studies in skin fibroblasts from patients suffering Parkinson’s Disease (PD) or Huntington’s Disease (HD) indicated that mitochondrial dysfunction in skin fibroblasts lead to decreased proliferation. Both studies showed a decrease in ATP production, but only the PD fibroblasts showed a depolarization of the mitochondrial membrane^36; 37^. The fibroblast responses seen in these studies had similarities to the phenotype described in chapter II and demonstrate levels of oxidative stress that lead to mitochondrial dysfunction that alter a fibroblast’s phenotype without inducing apoptosis.

Comparing the response to DOX exposure in cardiac fibroblasts to other injury models can be helpful, however the models mentioned did not also induce DNA damage. One study that compared fibroblasts of three different origins found that cardiac fibroblasts were resistant to etoposide-induced mitochondrial-mediated apoptosis. Even though the study used etoposide for cardiac injury, it did not report on DNA damage or p53 changes^38^. It was determined that constitutive expression of Bcl-2 was responsible for the blockade in cytochrome c release.

Despite the current field of research, a protective intervention has not yet been developed and proven effective in limiting chronic DOX-induced cardiotoxicity. The cardiac fibroblast, with its role in cardiac maturation and injury response, is a logical focus of research. The current work studies DOX-induced inhibition of mitophagy in cardiac fibroblasts that leads to inadequate stress responses and cellular dysfunction. The confirmation of a mechanism of damage and its amelioration indicates the significance of DOX’s effect on the cardiac fibroblast and the role it may play in the overall progression of DOX-induced cardiac disease.

## Material & Methods

### Animal Care & Protocol

C57BL6/J mice were purchased from The Jackson Laboratory (Bar Harbor, ME) and p53^+/-^ mice were a gift from the Lozano Lab at UT MD Anderson Cancer Center (Houston, TX). Animals were bred in-house to maintain the wild type C57BL/6J colony and to develop p53^-/-^ mice for cardiac fibroblast isolation. Mice housing rooms were equipped with temperature control and a 12-hour light-dark cycle. Mice were allowed water and standard chow *ad libitum*. Offspring were weaned at three weeks of age and separated according to sex. All mouse procedures were carried out in agreement with the Guide for the Care and Use of Laboratory Animals (National Research Council, National Academy Press, Washington DC, USA 2011) and were approved by the Institutional Animal Care and Use Committee at the University of Texas Health at San Antonio.

### Reagents

Antibodies purchased from Abcam (Cambridge, MA) were anti-p53 (ab26), anti-Parkin (ab15954), anti-nuclear lamin (ab16048), and anti-VDAC (ab15895). Anti-PGC1α was ordered from Santa Cruz (sc-15067) and anti-GAPDH was purchased from EMD Millipore (MAB374).

### Genotyping

Genotyping to determine p53^-/-^ mice was performed with the REDExtract-N-AMP Tissue PCR Kit (Sigma XNAT-100RXN). Ear punches were digested according to the kit protocol. DNA was amplified with the following primers:

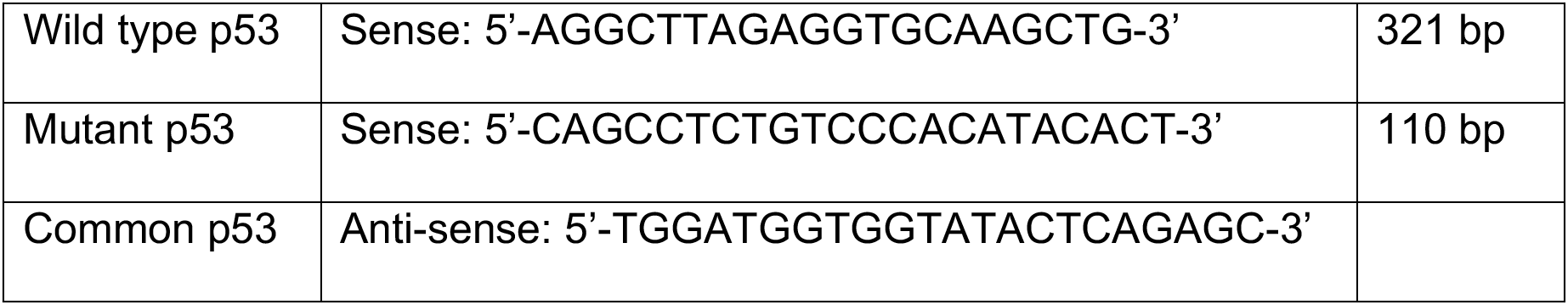

Samples were run on a 1.5% agarose gel (ThermoFisher J32802-22) in 1X TAE Buffer (EMD Millipore LSKMTAE50) with SYBR Safe DNA Gel stain (Invitrogen S33102) *Cell Isolation, Culture, and Treatment*

Cardiac fibroblasts (CFs) were isolated from the left ventricle of 6- to 8-week-old C57BL/6J and p53^-/-^ mice, both male and female. Mice were anesthetized with 3% isoflurane and euthanized by cervical dislocation. Excised hearts had the main vessels, atria, and right ventricle removed, then rinsed clean of blood in Hank’s Buffered Saline Solution (HBSS, Gibco 14190-136). After mincing with a scalpel, tissue was dissociated by mechanical disruption and three fifteen-minute collagenase II (Worthington LS00477) incubations. Cells were expanded in DMEM/F-12 50/50 media (Corning 16-405-CV) with 10% fetal bovine serum (Gibco 16000-044) and 1% Penicillin Streptomycin Solution (Corning 30-002-Cl).

For studies, CFs were plated and allowed to adhere overnight before treatment. Plating densities were calculated to achieve ∼80% confluence at adherence, except for growth assays. CFs were exposed for three hours to either 1, 3, or 5 µM DOX (TEVA NDC 0703-5043-01) or 3 µM FCCP (Abcam ab120081. After treatment, cells were washed two times with HBSS then covered in pre-warmed media and stored in the incubator. Assays were performed 3-72-hours after treatment. Variations in experimental format are noted where relevant in the methods below.

### Functional Studies

For proliferation studies, cells were plated in 96 well dishes (15,000 cells/mL) and images were obtained every 2 hours for three days. An Incucyte Live-Cell Analysis System (Essen Biosciences, Ann Arbor, MI) collected images and proprietary Incucyte software was used to assess confluence at each 2 hour time point. Migration assays were performed with QCM Chemotaxis Cell Migration Assay (EMD Millipore ECM508). Cells were treated with DOX or saline and then transferred to a Boyden chamber membrane in serum-free media after 24 hours. The lower chamber was filled with 10% FBS media. After 24 hours, migrated cells were stained and then the dye was extracted to measure absorbance. (EMD Millipore, Burlington, MA).

Three hours after treatment, cells were detached and a single cell suspension was generated. Cells were fixed with 80% ice cold ethanol in PBS. Cells were stained with propidium iodide dye for 25 minutes at 37°C. Fluorescence was quantified on a BD Calibur instrument by the UTHSCSA Flow Cytometry Core Center.

### Viability Assays

Cell mass after 24 hours was determined with a sulforhodamine B dye (SRB) assay. After treatment, cells were fixed, protein was stained with SRB, and then the dye was extracted. Absorbance was read and compared to absorbance at time zero. Membrane integrity at 3 and 34 after treatment was evaluated with a Trypan Blue Exclusion assay. Cells were incubated with trypan blue and then total cells and clear cells were counted. Cellular metabolism was tested via MTT reduction. Cells incubated in MTT solution and formazan crystals were solubilized with DMSO. Absorbance was read at 540 nm.

### Gene Expression Profiling

Exactly twenty-four hours after a three hour treatment of 3 µM DOX, total RNA was collected from cardiac fibroblasts utilizing the RNeasy Mini Kit. RNA was quantified using a Nanodrop ND-800, then 400 ng of total RNA was used in the RT^2^ First Strand Kit (Qiagen 330404) to convert to cDNA. Samples were prepared for Real Time qPCR with the RT^2^ SYBR Green qPCR Mastermix (Qiagen 330502) and run on the Applied Biosystems QuantStudio 7 Flex Real-Time PCR System. The threshold was manually set at 1.9 for every assay plate. Two pathway-directed RT^2^ Profiler PCR Arrays were selected from Qiagen: Mouse Extracellular Matrix & Adhesion Molecules (PAMM-013ZE-4) and Mouse Cytokines & Chemokines (PAMM-150ZE-4).

### Markers of Mitochondrial Health

Experiments for MitoSOX Red, Mitotracker Green (Fisher M36008 and M7514, respectively) were evaluated with microscopy and flow cytometry. Membrane potential was assessed with microscopy and a JC-1 Mitochondrial Membrane Potential Assay Kit (ab113850). Three hours after treatment, cells were incubated in the fluorescent Fisher dyes diluted in serum-free media. MitoSOX Red and Mitotracker Green samples incubated for 10 and 20 minutes, respectively. A single cell suspension in PBS was generated and samples were run on a BD LSR II flow cytometry instrument. Before treatment, cells were stained with 1 µM tetraethylbenimidazolycarbocyanin iodide (JC-1) for 10 minutes at 37 °C. The fluorescence was read three hours after treatment. To measure aggregate absorbance only, the excitation wavelength was set to 535 nm and the emission wavelength to 590 nm. To measure the monomer fluorescence, the excitation wavelength was set to 470 nm and the emission wavelength to 530 nm.

Microscopy images were obtained after dye incubation and 5 minutes of NucBlue (Molecular Probes R37605) staining. Images were obtained on a Nikon Eclipse TE2000-U scope.

### Cellular ROS Production

Cellular ROS production was measured using the DCFDA Cellular ROS Detection Assay Kit (Abcam ab113851) according to kit instructions. Cells were stained with 25 µM 2,7-dichlorofluorescin diacetate (DCFDA) in 1X buffer for 45 minutes at 37°C. The stain was removed, and cells were treated with 1, 5, or 10 µM DOX. A positive control consisted of cells treated with 50 µM tert-butyl hydrogen peroxide (TBHP, included in the kit). Negative and background controls included wells with media only, media and DCFDA stain, DOX-treated cells without stain, and cells without the DCFDA stain. Fluorescence was measured after three hours of treatment.

### Seahorse Energy Phenotype

Mitochondrial function was assessed with the Seahorse Energy Phenotype (Agilent Technologies 103275-100) according to the kit protocol. Briefly, cells are plated in a Seahorse XFp 96-well plate, with four wells containing media only. The sensor cartridge was hydrated with Seahorse XF Calibrant in a no-CO_2_ incubator. Cells were treated with 1, 3, or 5 µM DOX for three hours then washed with Seahorse Assay Media (10 mM glucose, 1 mM pyruvate, and 2 mM L-Glutamine) and placed in a non-CO_2_ incubator for one hour. A stock solution of 10 µM oligomycin and 10 µM FCCP was prepared and loaded into port A of the sensor cartridge to be dispensed during the assay.

The cartridge plate was then loaded into the Seahorse incubator and the Seahorse Phenotype Test Software was run. The Seahorse assay software obtains baseline measurements of oxygen consumption and acidification levels in the wells. After the oligomycin and FCCP are released into the wells, and measurements for oxygen and acidification are repeated. Optimal cell density, FCCP concentration, and oligomycin concentration were generated from a dose response titration of the three variables. Values were determined from Agilent recommended baseline levels of oxygen consumption and changes incited by stressor addition.

### Immunoblotting

For localization studies, cellular compartments were isolated using the Cell Fractionation Kit (Abcam ab109719). Fractionation was verified with GAPDH for cytosolic fractions, nuclear lamin for nuclear fractions, and VDAC for mitochondrial fractions. Protein content was measured with Pierce BCA Protein Assay (ThermoFisher Scientific 23225). Samples were run on 10% Bis-Tris Criterion Gels (BioRad, Hercules, CA) and transferred to nitrocellulose membranes. For loading control normalization, REVERT Total Protein Stain (Li-Cor 926-11010, Lincoln, NE) was used to acquire relative protein loaded for samples. Membranes were blocked in a 1:1 solution of Odyssey Blocking Buffer (Li-Cor 927-50000) and tris buffered saline. Fluorescent secondary’s from Li-Cor were used for visualization (925-32212 and 926-32211) along with a Li-Cor CLx imaging system. Densitometry measurements were acquired with Image J (NIH, Bethesda, MD).

### Immunofluorescent Cytometry

Cells were plated on coverslips in 12 well dishes. Three hours after treatment, cells fixed in 4% paraformaldehyde and permeabolized in 0.25% Triton X. Cells being probed for p53 and Parkin were blocked in 5% BSA after permeabolization and before incubating in primary antibody overnight at 4°C. After washing, cells were incubated in fluorescent secondary antibodies, stained with DAPI (Invitrogen D1306), and mounted with Prolong Gold Antifade reagent (Invitrogen P36930).

### Mitophagy Assay

Samples were prepared with the Dojindo Molecular Technologies, Inc (Rockville, MD) Mitophagy Detection Kit (MD01-10) according to manufacturer’s instructions. Briefly, cells were incubated in a mitochondrial-binding dye prior to treatment. Three hours after treatment ceased, cells were incubated with a lysosome dye. Samples were analyzed and cell populations were quantified on a BD LSR-II flow machine under the supervision of the UTHSCSA Flow Cytometry Core.

### Proximity Ligation Assay

Cells were plated on glass coverslips in a 12 well cell culture. Three hours after treatment, cells were fixed in 4% paraformaldehyde in PBS for 10 minutes at 37°C. Cells were permeabilized in 0.25% Triton X in PBS for 10 minutes at room temperature. The proximity ligation assay was performed using a Duolink In Situ Red Starter Kit Mouse/Rabbit (Sigma-Aldrich DUO92101). The cells were blocked with a solution provided by the kit and then incubated overnight at 4°C in a 1:500 dilution of primary antibodies for p53 and Parkin. Three negative control slides included replicates that did not incubate in the p53 antibody, the Parkin antibody, or either antibody. The antibodies were also diluted in a solution provided by the kit.

PLUS anti-rabbit and MINUS anti-mouse probes were diluted in the provided diluent and applied to the cells. Cells incubated with the probes for one hour at 37°C. The ligation solution was applied for thirty minutes at 37°C and then cells were incubated in the amplification buffer for 100 minutes at 37°C. Slides were mounted with a DAPI-containing mounting medium provided in the kit. Fluorescent images were taken at 20X magnification. At least ten fields of view per slide were obtained. Fluorescent signal was quantified using Image J and normalized to nuclear signal.

### Statistical Analysis

All statistical tests were performed using GraphPad Prism 7. A student’s T-test was used for comparisons between two groups, and a Welch’s correction for unequal variance was used when necessary. Comparisons between more than two groups were analyzed with a one-way ANOVA. A second order polynomial regression was fit to growth curves and comparisons of fit were conducted. Cell cycle distribution was analyzed with a Chi^2^ test. The threshold for the p value was set at 0.05, except for the gene array profiles where a more stringent threshold of 0.01 was used.

## Results

### Measures of Cardiac Function

Cardiac fibroblasts respond to injury by proliferating in and migrating to areas of damage. The ability to carry out these two functions are essential to cardiac fibroblasts’ role in development, as well as injury response. Therefore, the first items assessed when determining the effects of DOX on cardiac fibroblasts were growth and migration. Primary cardiac fibroblasts, isolated from C57BL/6J and p53^-/-^ mice were treated for three hours with 3.0 µM DOX, washed with Hank’s Buffered Saline Solution (HBSS), and placed in an Incucyte for live cell imaging.

**Figure 1** A, C shows DOX significantly reduced cardiac fibroblast growth (p<0.0001) in both cell strains. By 72 hours, the confluence of the treated cells was almost half that of untreated cells. After 36 hours, it appears that the reduction in confluence is due more to cell death than decreased proliferation. **Figure** 1B, D demonstrates the mask applied to measure cell confluence.

**Figure 1.**
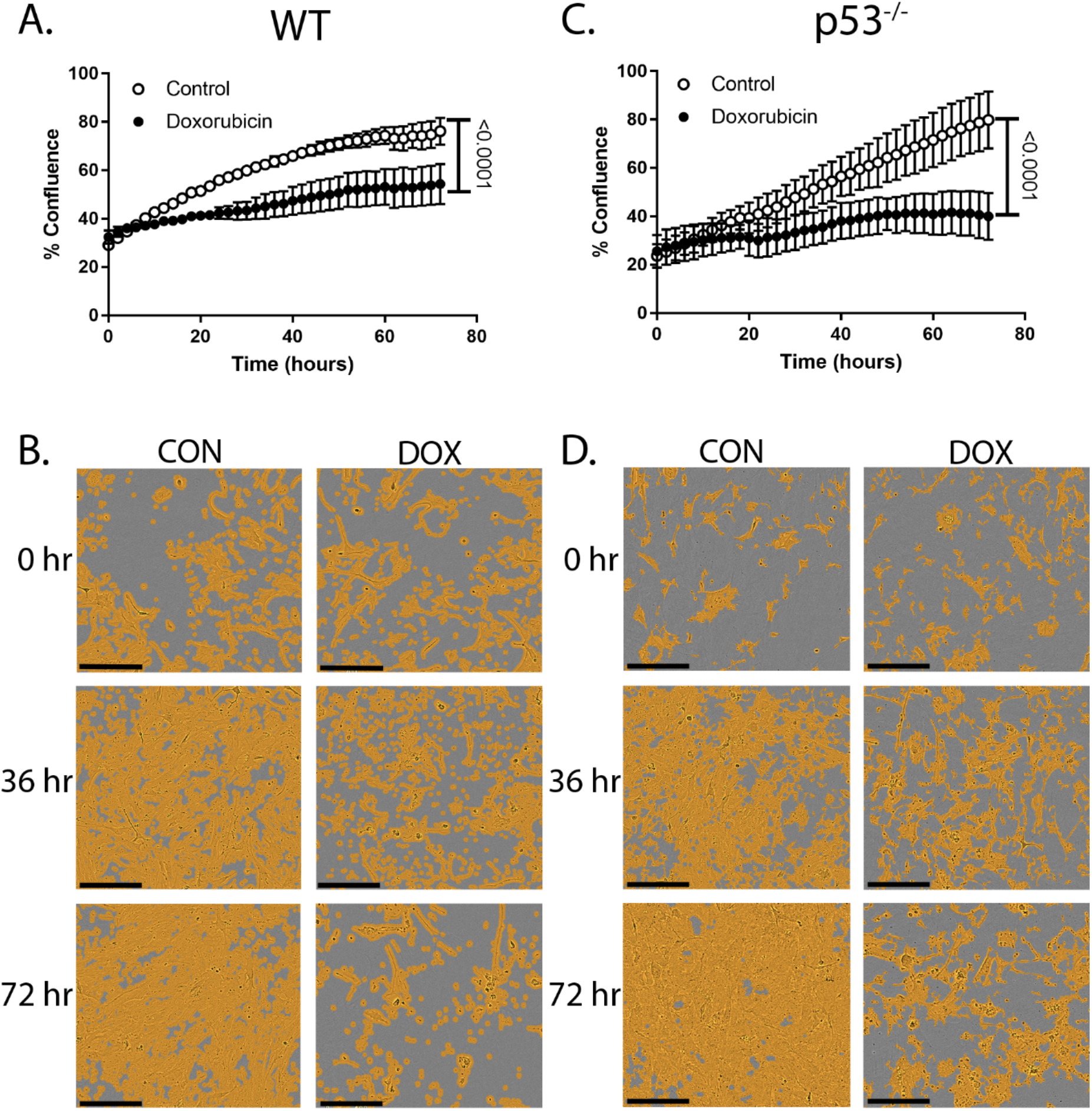
DOX Inhibits Proliferation. DOX inhibits proliferation in WT and p53^-/-^ cardiac fibroblasts (A, C). Cells were plated in a 96-well plate, treated for 3 hours, and then followed for 72 hours with live cell imaging. (B,D) are representative images of live cell imaging with the confluence mask. Graphs are an average of 3 biological replicates ± SEM. Scale bar is 200 µm.

Cell movement through a membrane in a modified Boyden Chamber assessed migration. DOX impeded migration by ∼15% in WT cells (p<0.0001), but p53 deletion restored the cell’s migratory ability (**Figure 2A**). After treatment and fixation with propidium iodide, cell cycle distribution was assessed via flow cytometry. WT cells showed a significant difference between the cell cycle distribution (p=0.001), indicating arrest at the G1/S checkpoint. There was no change in the percentage of cells in the G2 phase, with the complimentary increase/decrease seen in the G1/S phases, respectively. p53^-/-^ did not show the same cell cycle arrest (**Figure 2B, C**). The reduction in proliferation therefore has a different etiology in p53^-/-^ cells.

**Figure 2.**
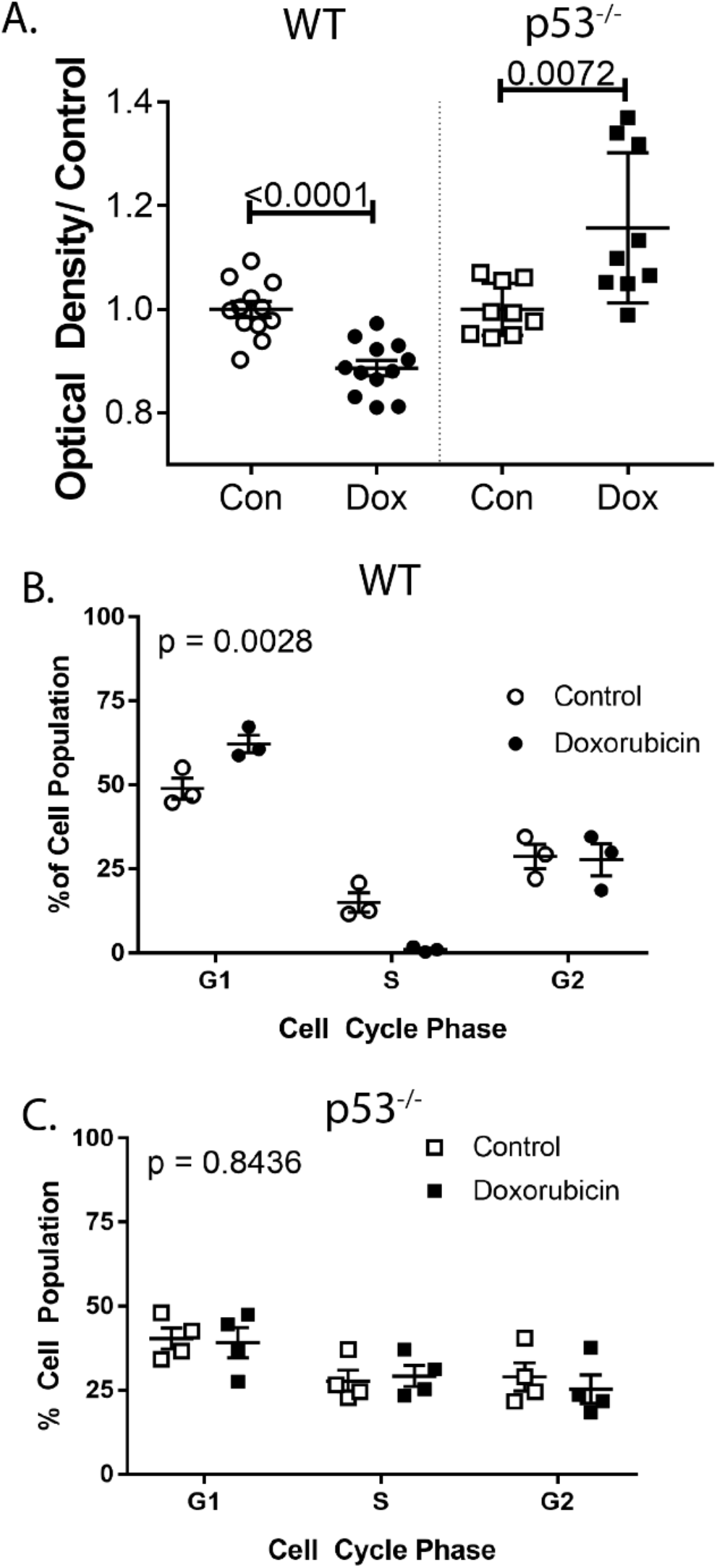
Migration is Restored in p53^-/-^ Cardiac Fibroblasts. While migration decreases relative to control in WT cells, it is increased in p53^-/-^ cells after DOX treatment (A). DOX exposure does induce cell cycle arrest at the G1/S checkpoint (B), but not in p53^-/-^ cells. Cell cycle distribution was analyzed with a chi^2^ test. Graphs are biological replicates ± SEM.

### Cell Viability after DOX Exposure

To determine if the reduced growth and migration were due to a decrease in proliferation, an increase in cell death, or some combination, viability assays evaluating membrane integrity, protein mass, and cellular metabolism were performed. Neither WT nor p53^-/-^ cells showed changes in cell viability at doses of 1, 3, or 5 µM at three or twenty-four hours post-treatment. In **Figure 3A, B**, it can be seen that 95% of cells were viable in all groups.

**Figure 3.**
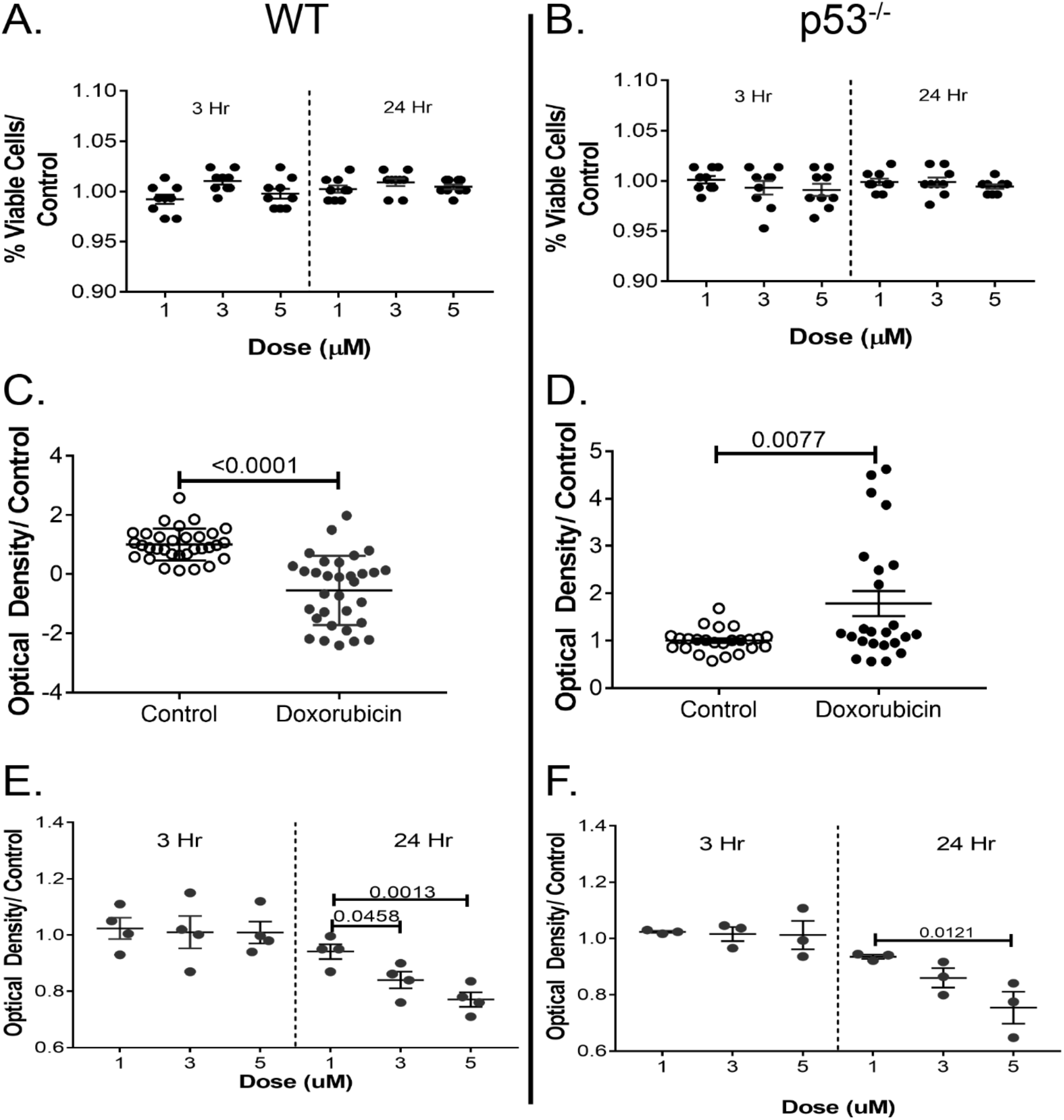
DOX alters metabolism, but not viability. Membrane integrity remained intact up to 24 hours after DOX doses up to 5 µM. Protein content decreased in WT cells exposed to DOX (C), but increased in p53^-/-^ cell after treatment (D). Both cell types showed an altered cellular metabolism at the 24-hour time point. Graphs are biological replicates ± SEM.

Reduction of the substrate MTT by oxidoreductase enzymes is an indication of cellular metabolic activity. MTT reduction remained constant compared to control at three hours post-treatment in all three doses. Twenty-four hours after treatment, metabolic activity decreased in a dose-dependent fashion (**Figure 3C**). The decreases at 3 and 5 µM were significant compared to both WT control and 1 µM DOX. For p53^-/-^ cells the reduction seen in the24-hour 5 µM group was statistically significant compared to the control and 1 µM groups, while the 3 µM group was only significant compared to control-a slight improvement from the WT studies (**Figure 3D**).

SRB was used to stain protein content in cells twenty-four hours after treatment with 3 µM DOX doxorubicin (**Figure 3E, F**). A significant decrease in protein content was seen after twenty-four hours in the WT treated cells (p<0.0001). The protein assay indicated an increase in protein production after DOX-exposure compared to control cells (p=0.0077).

### Gene Expression Profile

To determine how components of ECM remodeling and inflammatory signaling were affected by DOX, pathway-directed gene arrays assessed the expression of 168 relevant genes. Cells were harvested 24 hours after treatment and mRNA was isolated. After cDNA conversion, expression was measured using real time reverse transcriptase polymerase chain reaction (RT^2^-PCR).

Of the 168 genes, 50 were differentially expressed between the control and treated cells (**Table 1**). Twenty-eight genes were upregulated and twenty-two were downregulated in the treated cells. Structural genes, such as Col1a1, Col4a2, Col5a1, and Col6a1 were downregulated, while effectors of remodeling, such as MMPs 3, 8, and 12, were upregulated. An MMP inhibitor, TIMP3, was downregulated. Seven of the ten adhesion molecules with differential expression were downregulated in the DOX-treated cells. Sixteen inflammatory genes were significantly expressed, and all but two, were upregulated. Ccl2 and its receptor, Ccr4, were increased 40- and 50-fold respectively. Markers of cardiac dysfunction, Cxcl10 and Cxcl11, were increased 26- and 20-fold compared to control.

**Table 1.** Significant WT Gene Profiling Changes. 168 genes were measured, along with controls for genomic DNA contamination, polymerase activity, normalization. Significant gene changes were those that passed a limit of detection threshold, were two-fold greater or lesser than the control, and the p-value was <0.01. Red indicates an up-regulation in expression compared to control, while blue indicates a down-regulation. n=6

| ECM Structural |  | ECM Remodeling |  | Inflammatory Cytokines |  |  |  | Adhesion Molecules |  |
| --- | --- | --- | --- | --- | --- | --- | --- | --- | --- |
| Gene | Fold Change | Gene | Fold Change | Gene | Fold Change | Gene | Fold Change | Gene | Fold Change |
| Lamb3 | 5.4 | Mmp15 | 24.4 | Ccr4 | 53.0 | Csf1 | 7.0 | Cdh2 | 12.2 |
| Ecm1 | 2.3 | Mmp10 | 12.4 | Ccl2 | 42.5 | Ccl9 | 4.3 | Icam1 | 2.9 |
| Lama1 | -7.7 | Mmp8 | 7.5 | Cxcl10 | 26.4 | Il33 | 4.0 | Vcam1 | 2.5 |
| Col5a1 | -3.5 | Mmp13 | 4.6 | Cxcl11 | 20.5 | Il17b | 3.6 | Itgal | -3.7 |
| Col4a2 | -3.1 | Mmp12 | 3.4 | Osm | 18.1 | Nampt | 3.3 | Pcam1 | -3.0 |
| Lamc | -3.1 | Adamts8 | 3.3 | Ccl3 | 14.4 | Bmp2 | 3.0 | Ctnna2 | -3.0 |
| Emilin1 | -3.0 | Mmp3 | 3.0 | CD40l | 8.9 | Il4 | 3.0 | Ncam1 | -2.7 |
| Col6a1 | -2.9 | Adamts5 | -4.4 | Ccl24 | 8.4 | Cxcl5 | 2.1 | Itgax | -2.6 |
| Lama2 | -2.8 | Adamts1 | -3.6 | Il11 | -3.4 | Ccr8 | -2.1 | Itga4 | -2.3 |
| Col1a1 | -2.6 | Spp1 | -2.3 |  |  |  |  | Itgam | -2.1 |
| Fn1 | -2.3 | Timp3 | -2.2 |  |  |  |  |  |  |

Only 13 genes were differentially regulated between the control and treated p53^-/-^ cells (**Table 2**). Two genes, Hapln1 and Cdh3, were down-regulated, while the other 11 were upregulated. Most notable was the attenuation of the inflammatory genes. Only seven cytokines showed increased expression, and by only 4-fold or less. Ccr4, which was increased 50-fold in the WT cells, was only increased 3-fold in the p53^-/-^ cells. Ccl2, Cxcl10, and Cxcl11 expression was not significantly increased. Only three ECM remodeling genes were up-regulated while only one ECM structural gene was down-regulated.

**Table 2.**
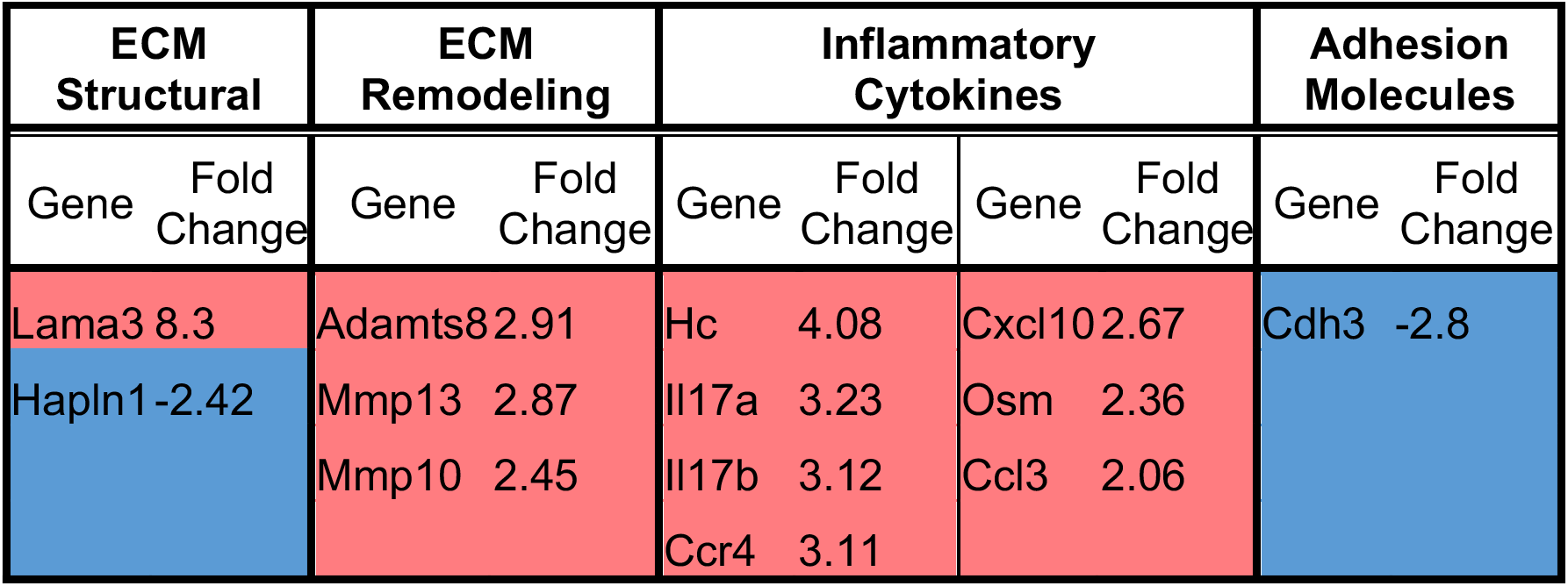
Gene Changes in p53^-/-^ Cells Exposed to Doxorubicin. 168 genes were measured, along with controls for genomic DNA contamination and polymerase activity. Significant gene changes were those that passed a limit of detection threshold, were two-fold greater or lesser than the control, and the p-value was <0.01.Red cells indicate an up-regulation in expression compared to control, while blue cells indicate a down-regulation. n=6

### Doxorubicin-Induced Oxidative Stress & Mitochondrial Health

In every cell where it has been studied, DOX induces the production of ROS. We determined that DOX does produce ROS in cardiac fibroblasts. Production of ROS was assessed by staining primary cardiac fibroblasts with a dye that fluoresces once oxidized. After staining, the cells were treated with multiple doses of DOX and the fluorescent signal read on a spectrometer. During treatment with DOX, ROS production increased in the cells in a dose-dependent manner (**Figure 4A**). At even a dose of 1 µM, there was a significant increase in ROS production during treatment.

**Figure 4.**
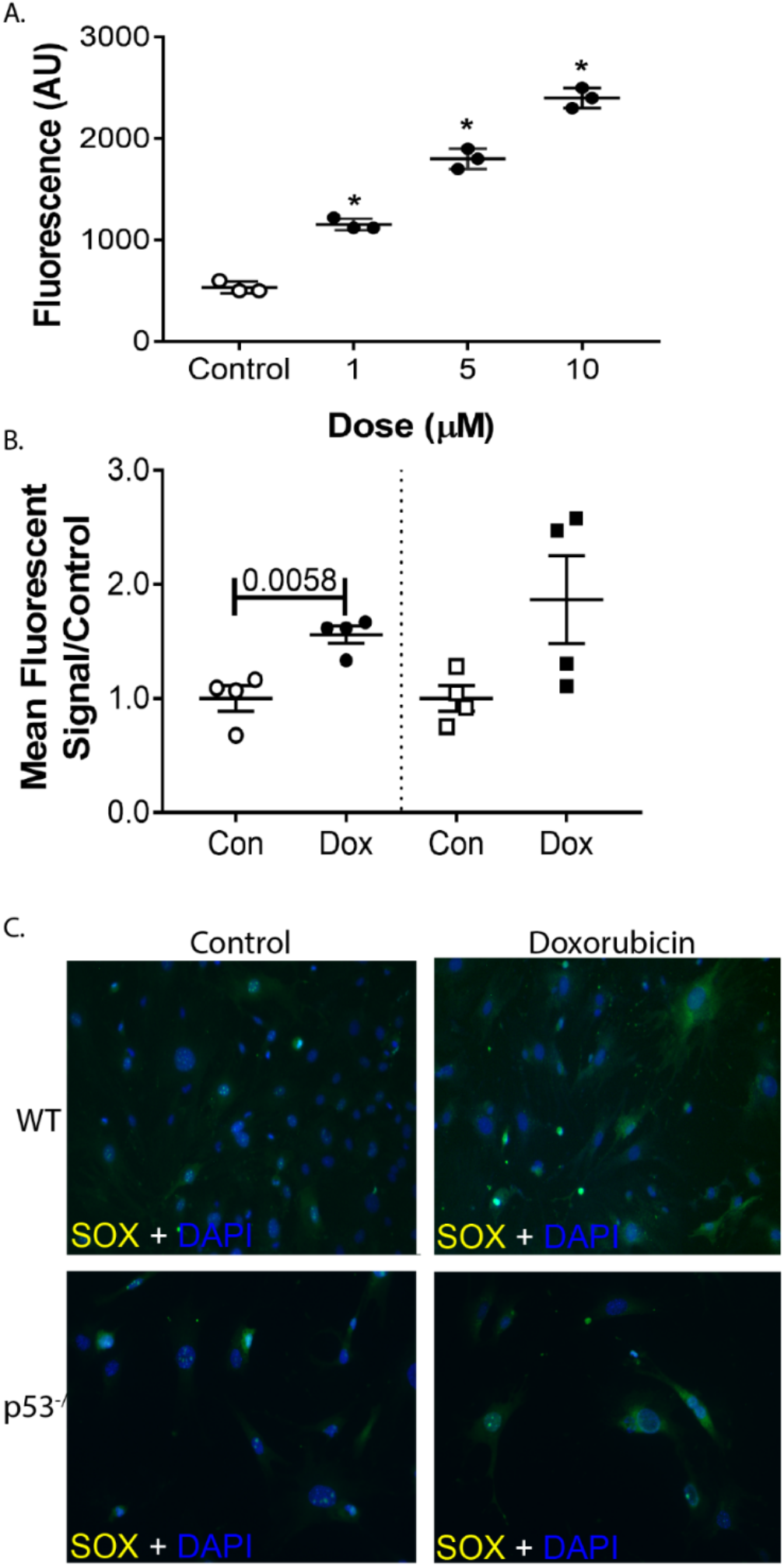
DOX induces cellular and mitochondrial ROS. DOX increased ROS production in a dose-dependent manner in the WT cell (A). WT cells demonstrate a significant increase relative to control cells in mitochondrial ROS (B). The response to mitochondrial ROS is more variable in the p53^-/-^ cells (B,C). Graphs are 3-4 biological replicates ± SEM.

To view mitochondrial-specific ROS, cells were plated on microscope slides and stained with MitoSOX Red after treatment. MitoSOX Red binds to mitochondria and fluoresces when oxidized. This fluorescence was quantified using flow cytometry. **Figure 4C** shows a representative staining of MitoSOX Red in control and treated cells, along with quantification of relative mitochondrial ROS. DOX did increase the presence of ROS at the WT mitochondria compared to control (p=0.0058). This was not true for p53^-/-^ cells.

Mitochondrial health after DOX exposure was assessed with two markers. The dyes bind to mitochondria and relate the relative mitochondrial load (Mitotracker Green) and mitochondrial membrane potential (JC-1). Mitochondrial mass in primary cardiac fibroblasts was assessed qualitatively by fluorescent microscopy and quantitatively by flow cytometry. Mitochondrial mass in the either cardiac fibroblasts strain was not changed by DOX exposure (**Figure 5A, B**). If mitochondria have a healthy polarized membrane, the JC-1 dye will aggregate in the mitochondrial membrane. As the membrane depolarizes, the dye diffuses into the cytoplasm as monomers. There is also a shift in the excitation/emission spectra of the dye. JC-1 aggregates fluoresce in a red wavelength, while the monomers fluoresce in a green wavelength. The intensity of the fluorescent signals can be compared to give a ration of healthy to unhealthy mitochondria in a sample. Again, membrane potential in primary cardiac fibroblasts was visualized with fluorescent microscopy. The fluorescent signal was quantified on a spectrophotometer. DOX decreased the average mitochondrial membrane potential, indicating depolarization of the membrane and a lower ration of healthy to unhealthy mitochondria compared to control samples (**Figure 5C, D**). The change was ameliorated in the p53^-/-^ cells.

**Figure 5.**
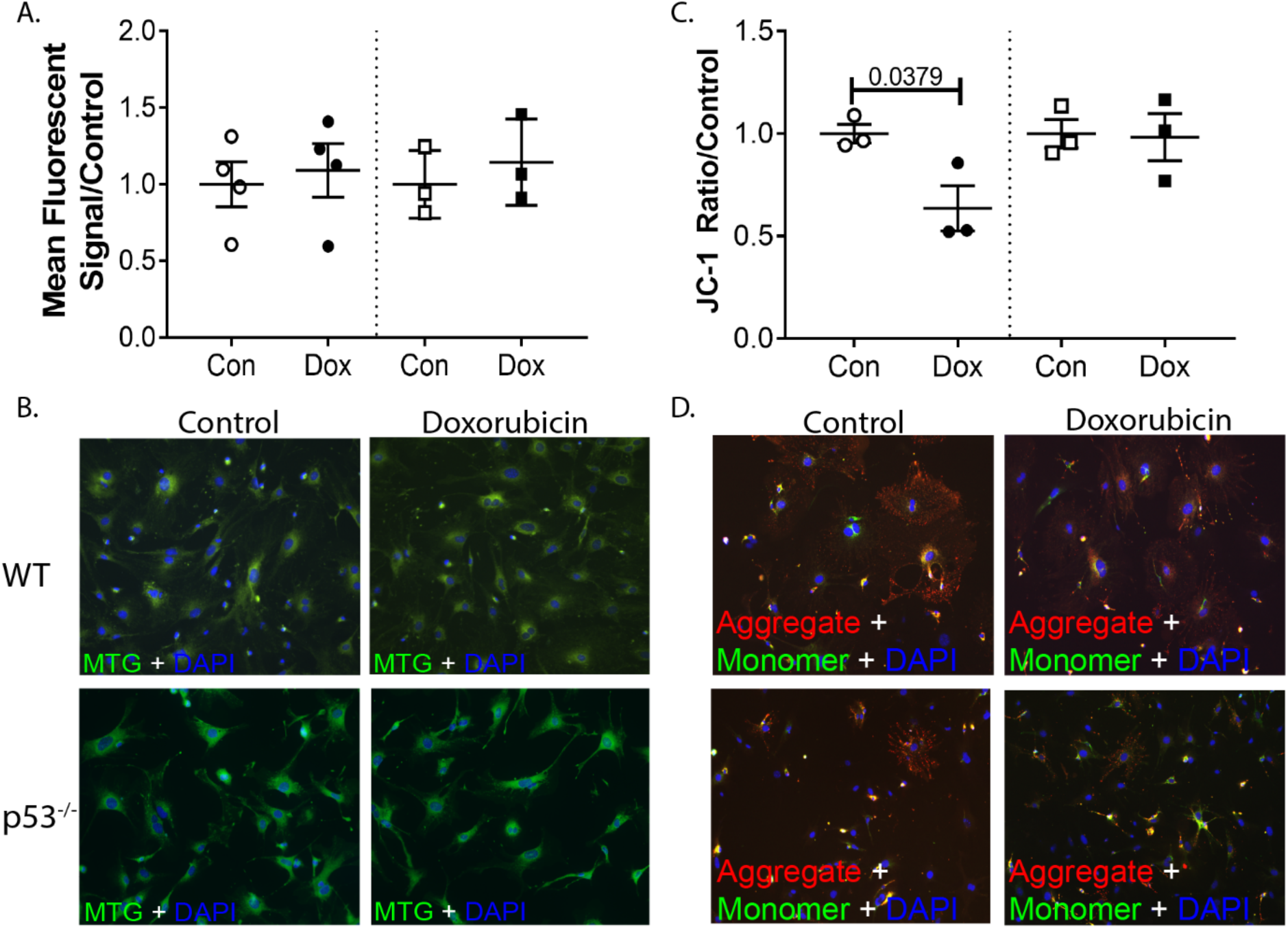
p53^-/-^ attenuates the DOX-induced mitochondrial depolarization. DOX has no effect on net mitochondrial mass (A, B), but does induce mitochondrial membrane depolarization in WT Cells (C, D). Panel D demonstrates the monomer vs aggregate staining of mitochondrial membranes Graphs are a combination of three biological replicates ± SEM.

Next, mitochondrial function was assessed to determine if the increased oxidative stress and membrane depolarization caused mitochondrial dysfunction. Oxygen consumption (OCR) and extracellular acidification (ECAR) rates were obtained at baseline and after mitochondrial stressors with the Seahorse Energy Phenotype Assay. At baseline there was a dose-dependent decrease in WT OCR that was not seen in the ECAR baseline measurements. Despite the lower baseline OCR, there was not a statistically significant difference in percent OCR change (stressed OCR/baseline OCR) among the control and treated groups (**Figure 6A, C**). There was a shift towards glycolysis in the treated cells in response to the mitochondrial stressors. The percent change was statistically significant in the 3 and 5 µM groups (**Figure 6E, G**). p53^-/-^ cells showed a similar dose-dependent decrease in OCR at baseline compared to control (**Figure 6B, D**). After stress, the DOX-treated cells were able to increase mitochondrial respiration similar to control cells. At the 3 and 5 µM doses, samples actually showed a slight increase in metabolic potential compared to control cells (p = 0.0053 and 0.0002, respectively). The increased metabolic potential indicates that DOX exposure primed the cells for increased mitochondrial respiration once under stress. The glycolytic shift seen in WT cells compared to control cells after stress was not seen in the p53^-/-^ cells (**Figure 6F, H**).

**Figure 6.**
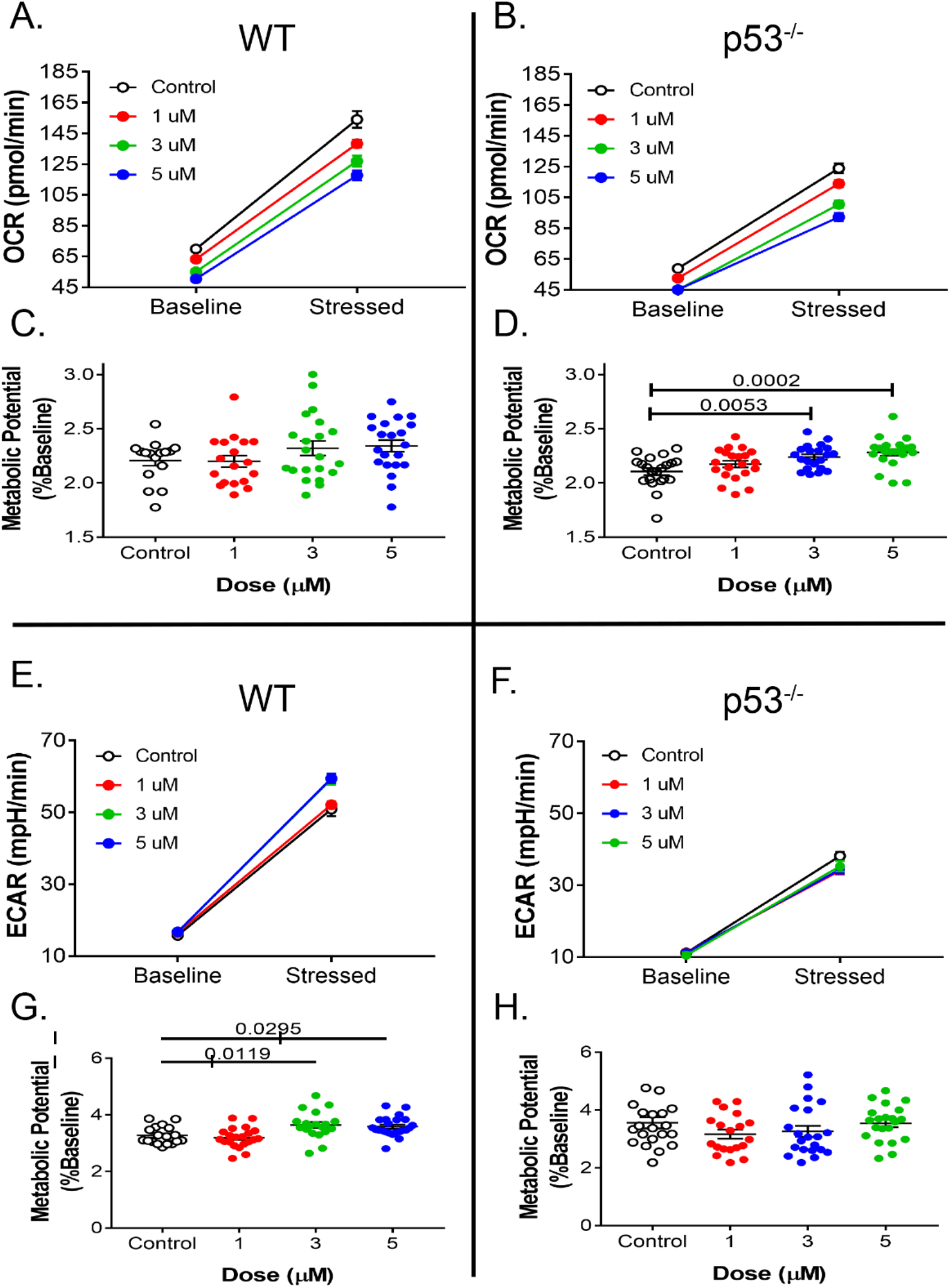
DOX-induced mitochondrial dysfunction is ameliorated in p53^-/-^ fibroblasts. In response to stress, WT cells increase glycolysis (E, G) as opposed to mitochondrial respiration (A, C) after DOX exposure. p53^-/-^ cells exposed to DOX were able to respond to the stressors with an increase in mitochondrial respiration beyond the control cells.

### DOX Induces Mitophagy in Cardiac Fibroblasts

To test the cell’s ability to undergo mitophagy in response to DOX, it must first be shown that the DOX-induced mitochondrial damage is sufficient to prompt mitophagy. To assess mitophagy in primary cardiac fibroblasts, we used a dual staining technique. **Figure 7** demonstrates the two fluorescent dyes used to quantify lysosome content, mitophagy, and the overlap of these two components. DOX exposure does increase the size and number of lysosomes compared to control (left column). There is only some overlap between the lysosomal dye and the mitophagy dye in the DOX samples. It is evident in the FCCP-treated sample that the two dyes co-localize with minimal lysosome-only dye visible. Increased lysosomes and Parkin expression (discussed later) indicate that mitophagy is induced in DOX- and FCCP-treated cells

**Figure 7.**
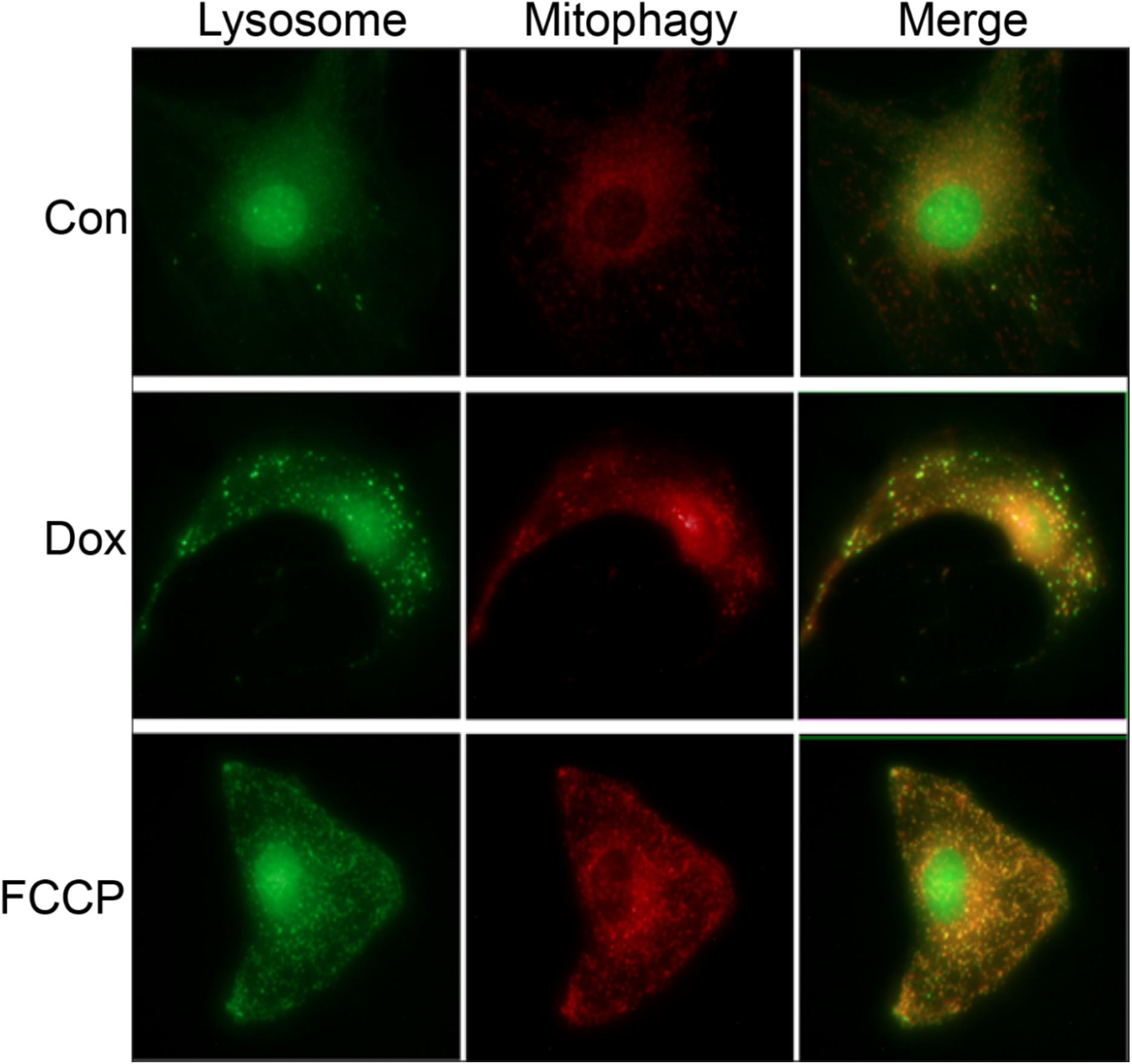
Representative Images of lysosome/mitophagy staining in WT cells. Cells were stained with a mitochondrial dye before treatment and a lysosomal dye after treatment. Fluorescent intensity of the mitochondrial dye increases when it is exposed to an acidic pH.

After confirming that mitophagy is induced in response to DOX-induced mitochondrial damage, it was necessary to assess whether or not the cells were able to complete the process of mitophagy. This was assessed by measuring the relative amount of overlap between the mitochondrial marker and lysosomal markers via flow cytometry. Before treatment with DOX or FCCP, cells were stained with a mitochondrial-binding dye. Under normal pH conditions, the mitochondrial dye exhibits fluorescence. However, in an acidic environment, as when an autophagosome and lysosome fuse, the fluorescent signal of the dye will increase. The shift in fluorescent signal, along with co-localization of the dyes indicates the final step of mitophagy. To quantify this, separate samples were prepared for flow cytometry. The population of cells with high intensity signal for both wavelengths were considered “double positive” and indicative of an active mitophagy process. This quantification can be seen in **Figure 8A**. There was almost no difference between the control and DOX-treated samples in the percentage of double positive cells. However, there was an increase in double positive cells in the FCCP-treated fibroblasts compared to control and DOX samples (p=0.0319). Mitophagy in p53^-/-^ was difficult to assess. This is possibly due to the unopposed activity of Parkin. PGC1α was similar between groups in both cell strains (**Figure 8B**).

**Figure 8.**
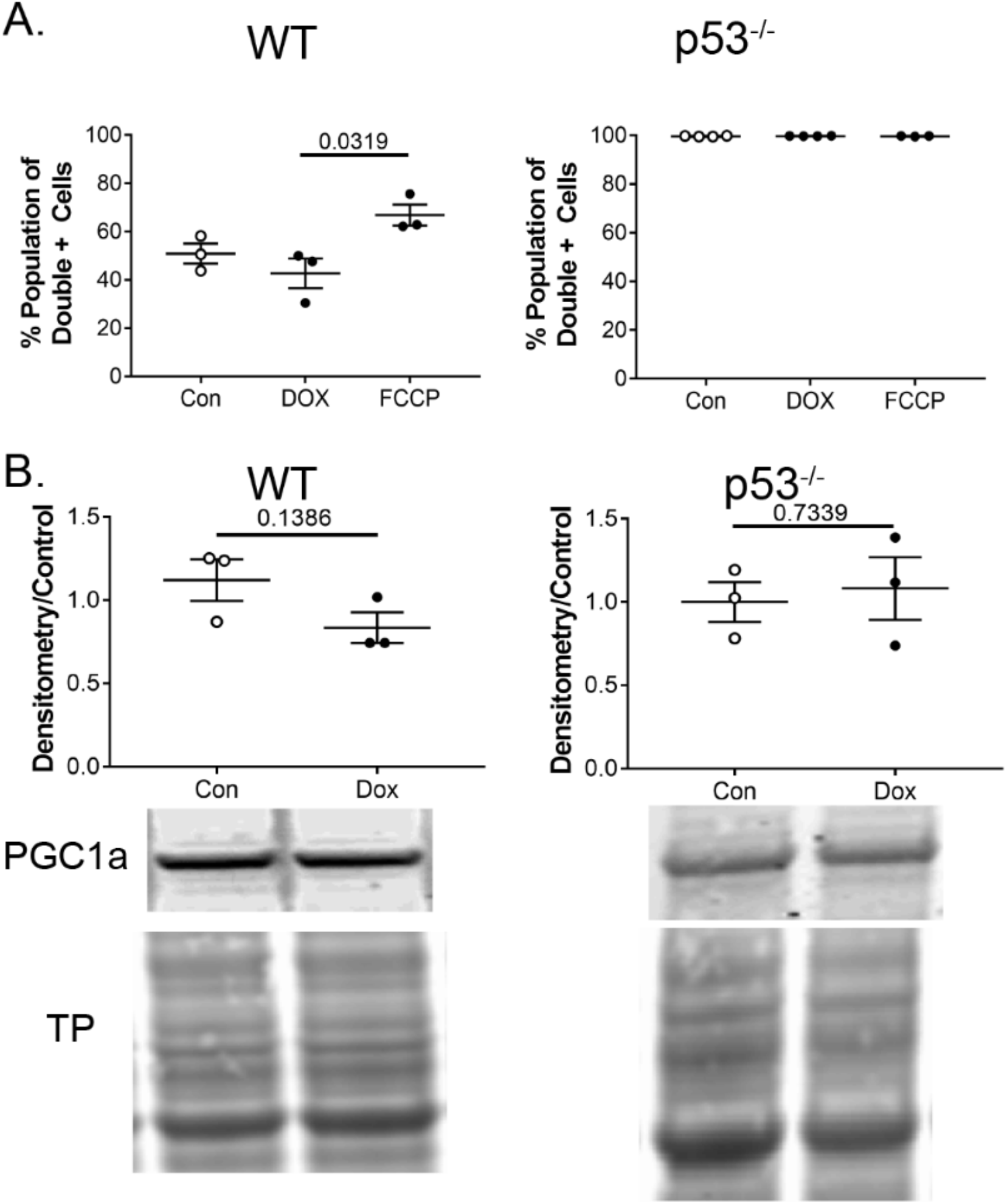
Mitophagy and biogenesis balance. (A) DOX-exposed WT cells were unable to respond to mitochondrial stress in a manner similar to FCCP-treated cells (left panel). Control cells in p53^-/-^ indicated high baseline levels of mitophagy (right panel). (B) PGC1α, a regulator of biogenesis, did not increase in either cell type in response to DOX.

### p53 Prevents Parkin Localization

To understand the role of p53 in mitophagy, it was confirmed that p53 expression in primary cardiac fibroblasts was increased after DOX exposure (**Figure 9A**). Not only was DOX increased almost three-fold compared to controls, it was increased by about the same amount compared to FCCP-treated samples. This validates the use of FCCP as a positive control for mitophagy without the interference of DOX-induced p53 upregulation. Panel B of **Figure 9** confirms the use of p53^-/-^ mice that have a mutated p53 gene. The mutated gene has exons 2-6 deleted, which includes the promoter region of the gene. These mice do not produce p53 and will begin developing tumors around four months of age, a time point after we isolate cardiac fibroblasts.

**Figure 9.**
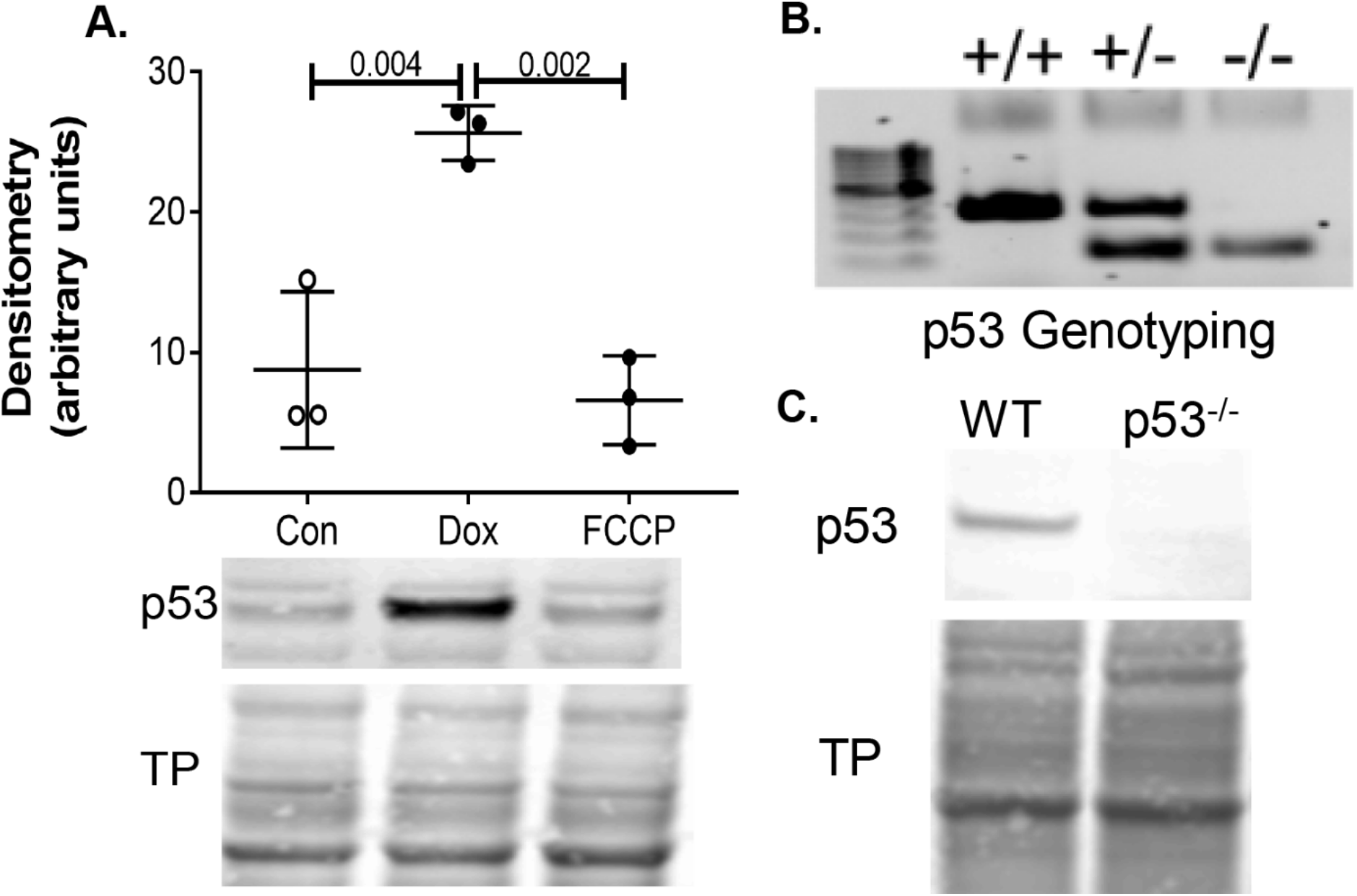
DOX upregulates p53 expression in WT cells. (A) DOX treatment of cardiac fibroblasts increases p53 expression compared to control and FCCP samples. Homozygous p53 knockout mice (B) do not express p53 (C).

**Figure 10** demonstrates the staining patterns of Parkin and p53 seen in control, DOX-, and FCCP-treated cells. In control cells, p53 remains diffuse throughout the cytoplasm. Parkin expression is not upregulated as it is with agents that induce mitochondrial dysfunction. Treatment with FCCP shows a distinct upregulation of Parkin and possibly a small increase of p53. While p53 remains diffuse throughout the cell, the Parkin staining has a linear pattern, possibly because it is localizing to the mitochondria. In the DOX-treated cells there is an upregulation of p53 and much of it enters the nucleus. However, there is still cytosolic p53. Parkin is upregulated but remains diffuse throughout the cell with only a couple areas where it might demonstrate a linear pattern. The most notable finding in the p53^-/-^ cells was the overexpression of the unopposed Parkin. Exposure time had to be radically reduced to avoid oversaturation, (5 s in WT samples, compared to 500 ms in p53^-/-^). Increased Parkin can be seen in control cells.

**Figure 10.**
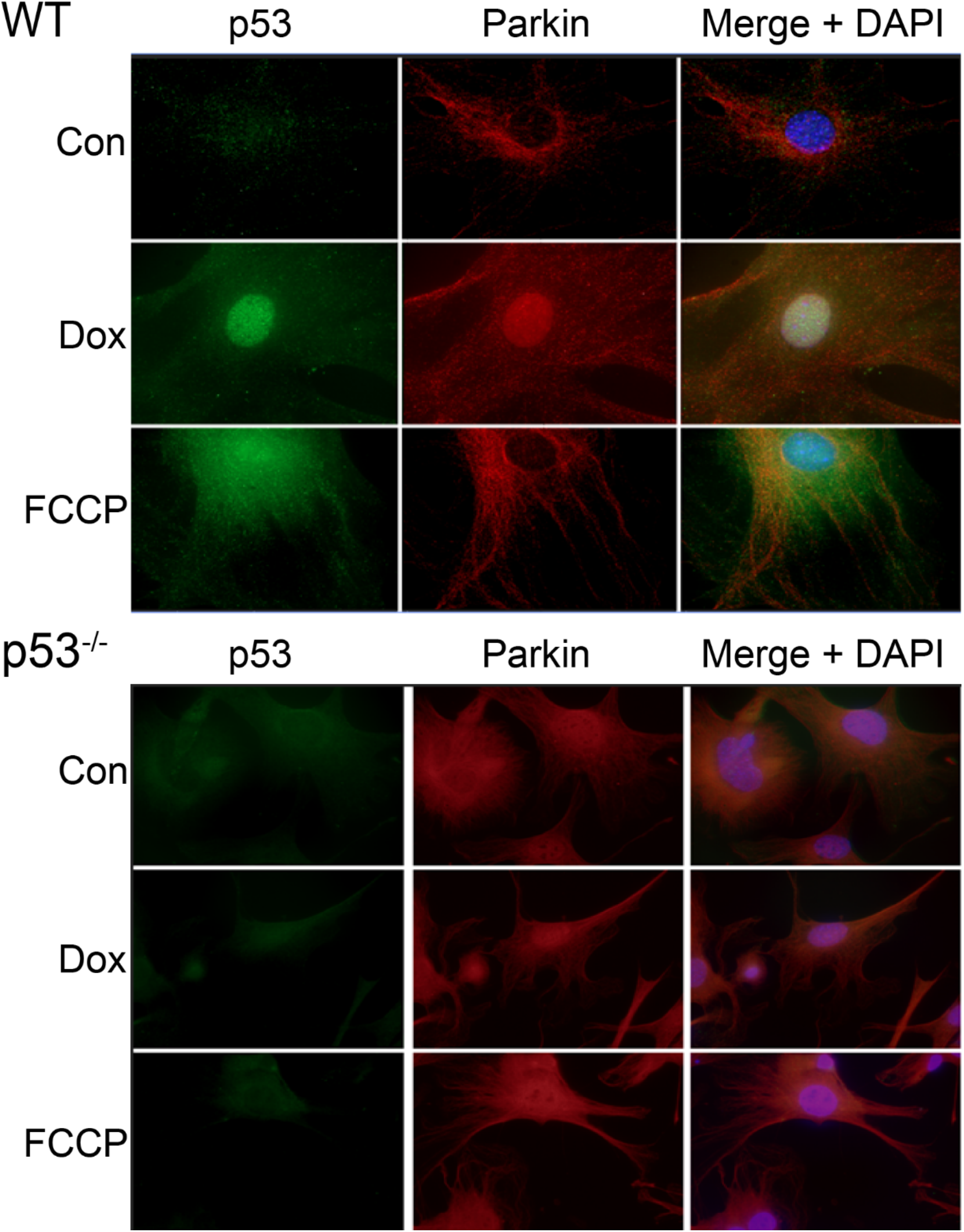
Parkin and p53 exhibit different staining patterns in response to FCCP and DOX. DOX and FCCP exposure upregulate Parkin expression in WT cells, but Parkin remains diffuse in the DOX-treated cells. Parkin expression in p53^-/-^ cells was upregulated and exposure time had to be decreased compared to WT cells.

To determine if the p53:Parkin interaction was the cause of the reduced mitophagy in DOX-treated cells, two items were assessed. The first was to verify if Parkin was able to translocate to the mitochondria after DOX treatment. The second item was to establish the interaction between p53 and Parkin. After treatment with DOX or FCCP, cell samples were processed to yield a cytoplasmic fraction and a mitochondrial fraction. Immunoblotting with anti-Parkin showed that Parkin was present in the cytoplasm of the control, DOX, and FCCP samples. However, minimal Parkin was present in the mitochondrial fractions of the control and DOX samples (**Figure 11B**). The Parkin present in the mitochondrial fraction of the FCCP sample had a molecular weight shift up, indicating a slight increase in the molecular weight. In order for Parkin to locate to the mitochondria, it must first be ubiquinated and once it reaches the mitochondria it is phosphorylated by Pink1. This finding agrees with similar studies that demonstrate an increase in the molecular weight of mitochondrial-associated Parkin due to monoubiquitination^39^. p53^-/-^ cells did show an increase in Parkin in either the DOX or FCCP groups (**Figure 11C**). However, this appears to be due to a higher basal level of Parkin in the control samples. This may be due to the lack of Parkin regulation in these cells. Obtaining mitochondrial fractions with the shifted Parkin was difficult, possibly due to early degradation. Even with various protease inhibitors, samples could often only be thawed once, or there was the risk of protein degradation. However, Parkin with the molecular weight shift was observed in the mitochondrial fraction of the p53^-/-^ cells. Quantification was not possible without at least three samples for the mitochondrial fraction.

**Figure 11.**
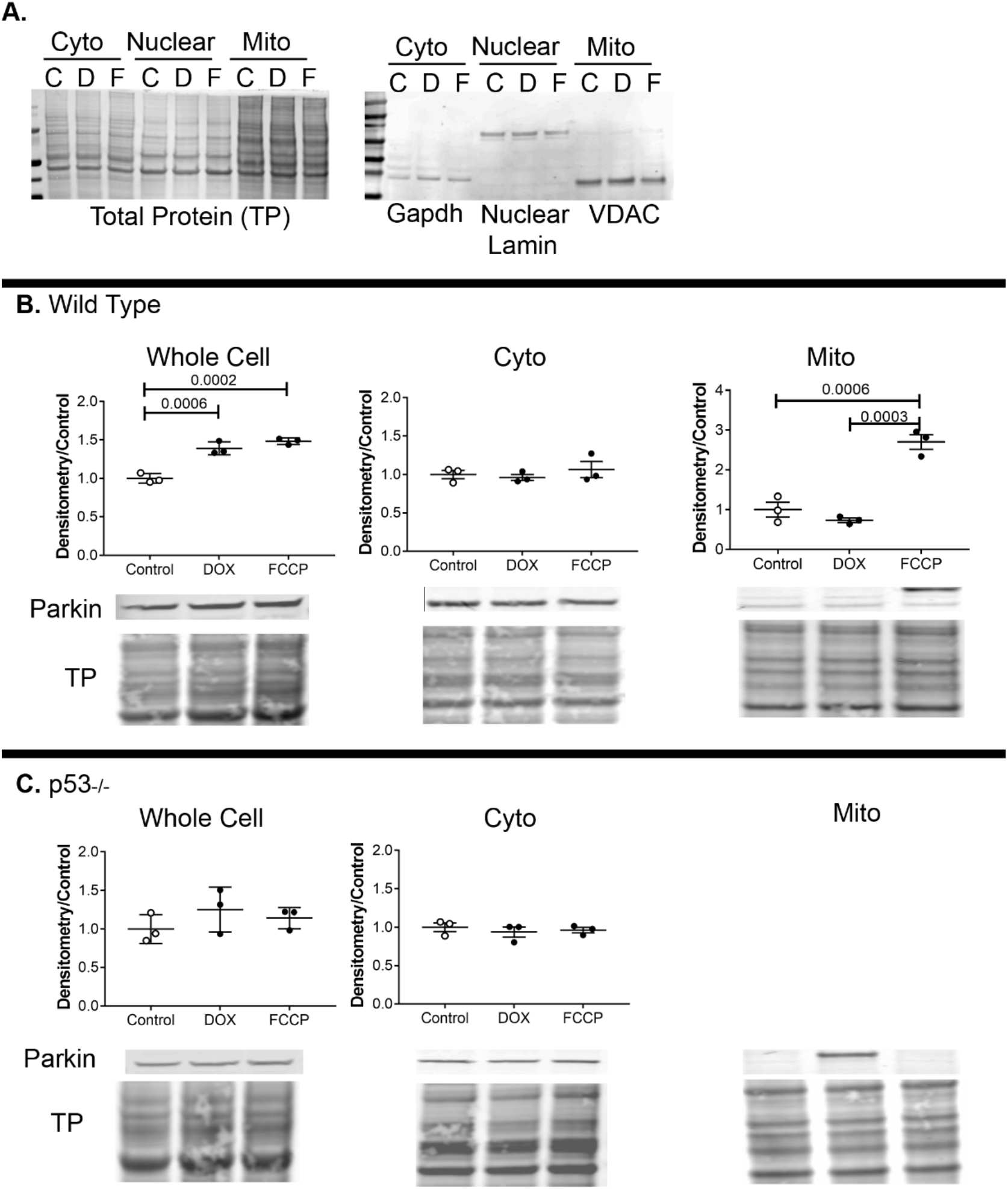
Parkin cannot localize to damaged mitochondria after DOX exposure. (A) Fraction efficiency and specificity confirmed with subcellular specific antibodies. (B) Parkin expression increased after DOX and FCCP exposure compared to control in WT cells. (C) Parkin did not increase in whole cell lysates of p53^-/-^ cells. While unable to quantify due to an n of 1, Parkin was still observed in the mitochondrial fraction after DOX exposure.

To determine if p53 was binding to Parkin, we used a proximity ligation assay. Fixed primary cardiac fibroblasts were probed with anti-p53 and anti-Parkin antibodies. PLUS and MINUS probes capable of forming circular DNA constructs were pre-conjugated to the secondary antibodies. Samples were incubated with ligase to create the circular DNA. Lastly, the circular oligonucleotides are amplified with fluorescent labels. Increased fluorescent puncta indicates an increase in the p53:Parkin interaction. Proteins must be within 40 nm of each other for ligation of the circular DNA to occur. DOX-treated cells showed a greater number of fluorescent puncta and increased fluorescent intensity compared to control cells (**Figure 12B, C**). Negative controls did show a small amount of non-specific signal in the DOX samples processed with anti-Parkin antibody, but no anti-p53 primary antibody. In p53^-/-^ fibroblasts there was an increase in the background of these same negative controls. Again, this is most likely due to the overexpression of Parkin in these cells.

**Figure 12.**
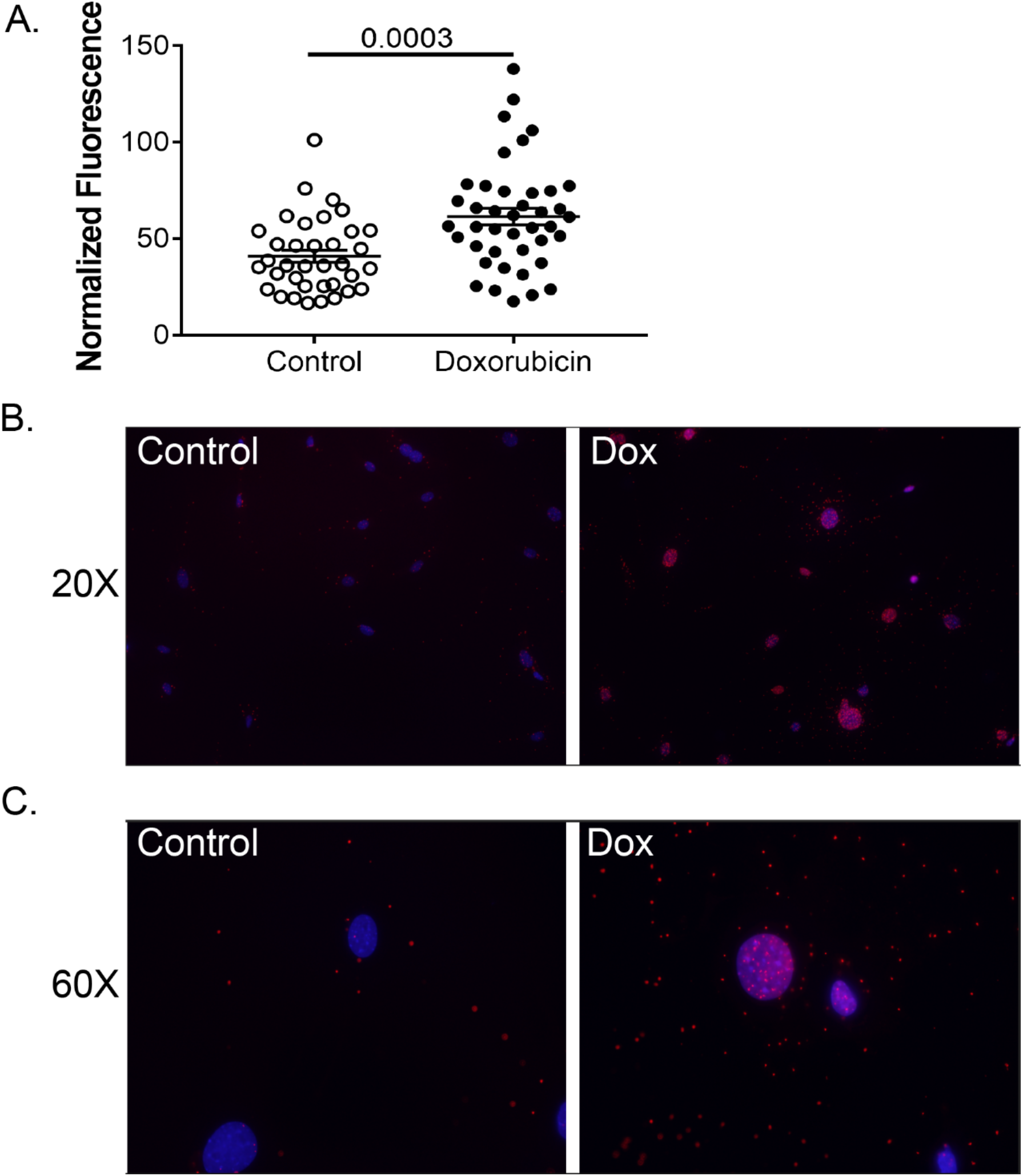
DOX increases Parkin:p53 interactions. (B, C) A proximity ligation assay demonstrates the increased interactions between the Parkin and p53 proteins in WT cells exposed to DOX. (A) Quantification of fluorescent signal. Quantification consists of 3-4 biological replicates with at least 10 fields of view acquired at 20X magnification. Graph is technical and biological replicates ± SEM.

**Figure 13.**
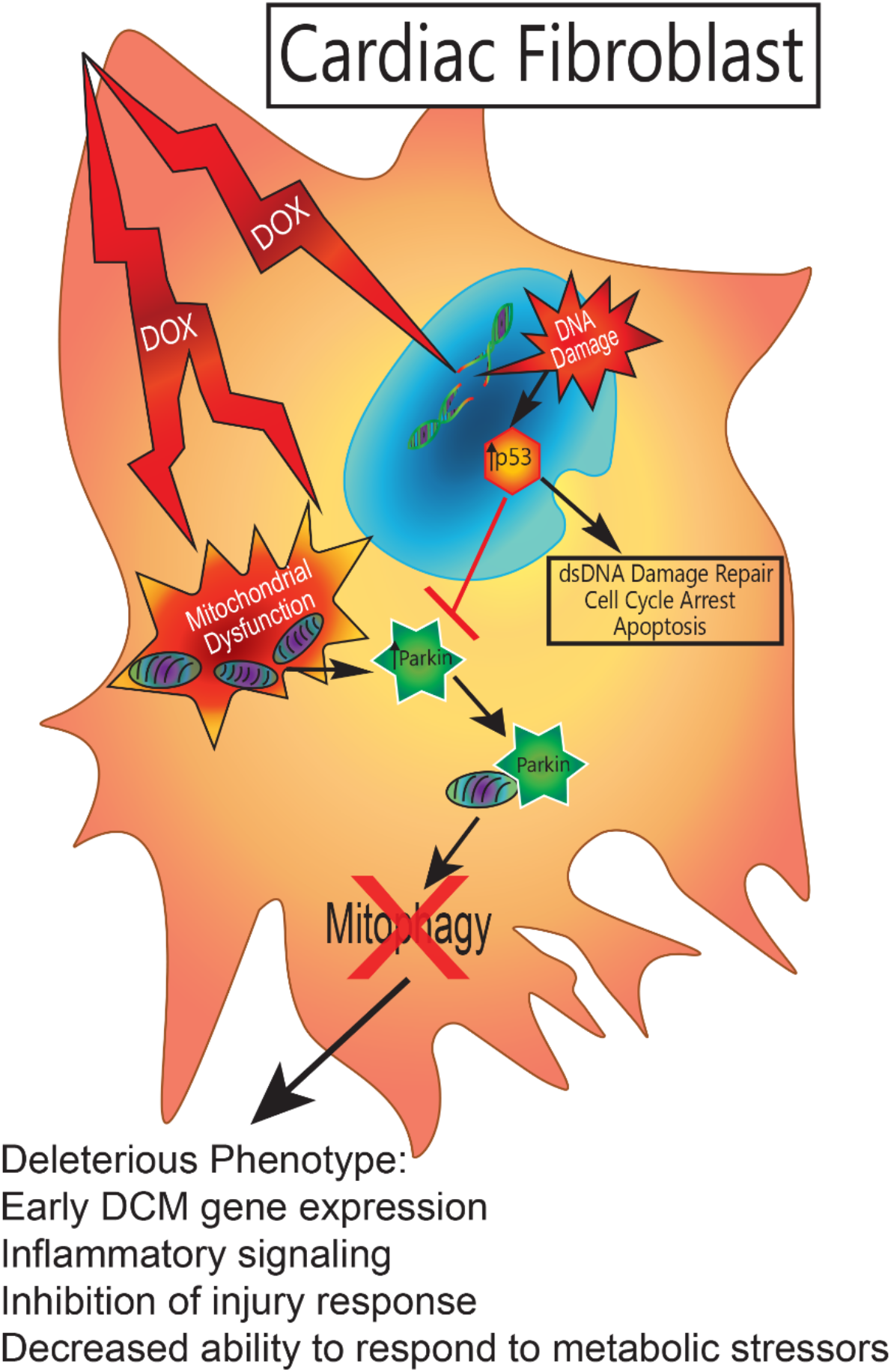
Parkin/p53 Interference. DOX induced a deleterious phenotype in WT cardiac fibroblasts. Due to the cell’s inability to respond to multiple stressors, an early DCM gene profile was adopted. This included increased cardiac remodeling genes and increased inflammatory signaling. The phenotype also removes the cell’s ability to respond to injury by inhibiting proliferation and migration. Lastly, the cell is open to increased damage as it is unable to respond to metabolic stressors due mitochondrial dysfunction and deficits in clearing the dysfunctional mitochondria.

## DISCUSSION

Healthy cardiac fibroblasts are essential for normal cardiac performance and the ability to respond to stress. While integral to directing the response to injuries, such as myocardial infarctions, cardiac fibroblasts are also essential for proper electrical conduction in the heart as they regulate calcium homeostasis through myocyte:fibroblast coupling^40; 41^. In addition, cardiac fibroblasts help maintain tissue structure by regulating the deposition and remodeling of collagen and other ECM components ^42; 43^. Recently, the novel concept of “mitoception” has garnered attention. This concept involves the transfer of mitochondria from one cell to another as a way of maintaining healthy mitochondrial function^44^. Considering the functions of cardiac fibroblasts in normal and pathologic states, understanding their role in DOX cardiotoxicity is crucial to developing a preventive or therapeutic treatment. The information generated in this body of work provides evidence for one mechanism of DOX damage that centers on the cardiac fibroblast and the significance of the cardiac fibroblast in the progression of DOX cardiotoxicity.

The DOX-induced phenotype observed in this study did not indicate an activation to a myofibroblast. There were similarities to a senescent phenotype, such as arrested proliferation and increased cytokine production. It is likely that a subgroup of the population did enter senescence, however the cellular morphology of the cells does not support this phenotype for the majority of the culture, WT or p53^-/-^.

In response to DOX, cells throughout the body upregulate expression of the tumor suppressor, p53^33; 45; 46^. In the heart, this occurs in cardiac myocytes, and as our data shows, in cardiac fibroblasts. In response to DNA damage, p53 undergoes a translocation to the nucleus to initiate cell cycle arrest, DNA damage repair, or apoptosis. In our model, p53 causes cell cycle arrest that generally does not progress to apoptosis in the timespan studied. *Zhan et al*, demonstrated an increase in ataxia telangiectasia mutated (ATM) protein after DOX exposure, indicating a DNA damage response. *Zhan et al*. also noted that this increase was predominantly in cardiac fibroblasts, as opposed to other cells of the heart such as cardiac myocytes^29^. This data suggests that cardiac myocytes do not suffer DNA damage to the same extent as cardiac fibroblasts, further supporting the idea that DNA damage and mitochondrial damage are both significant stresses in the fibroblast, as opposed to the myocyte or cancer cell. From immunocytochemistry, it is evident that an appreciable amount of p53 remains in the cytosol. *Hoshino et al.* described the direct interaction between cytosolic p53 and Parkin that prevented Parkin from binding at the mitochondrial membrane^30^. They also demonstrated that this interaction was sufficient to disrupt the process of mitophagy in cardiac myocytes. In fact, whole-body p53 depletion in mouse models attenuated cardiac dysfunction and decreased fibrosis after DOX exposure compared to wild type mice^33^. When p53 deletion was restricted to cardiac myocytes, the same benefits were not seen^34^.

There are several reasons why DOX affects myocytes differently than tumor cells. The cytotoxic effect of DOX on tumor cells relies on inducing apoptosis via DNA damage during cell replication. However, cardiac myocytes are terminally differentiated. In terms of metabolism, cardiac myocytes are dependent on oxidative phosphorylation to meet the high energy demands of the cell, while tumor cells switch to glycolysis to meet their energy needs even in an aerobic environment. These differences in energy sources and proliferation dictate the differential effects of DOX on these cell types, but offer no concise explanation for the effect of DOX on cardiac fibroblasts. Quiescent cardiac fibroblasts do not have the same oxygen and energy demands as myocytes, but cardiac fibroblasts do produce most of their ATP through cellular respiration. In terms of replication, cardiac fibroblasts may not replicate as often as tumor cells, but they are not terminally differentiated like myocytes and replicate throughout their lifetime. These factors suggest that DOX may elicit a class of pathological effects on cardiac fibroblasts that does not resemble what is seen in either myocytes or tumor cells.

Mitophagy at homeostasis is generally thought to be mediated by BNIP3. However, studies have shown that both tumor cells and cardiac myocytes can utilize the BNIP3 pathway under stress^47; 48^. In the absence of Parkin, these cell types may be able to maintain mitophagy under stress. The role of BNIP3 in relation to cardiac fibroblasts is not well studied. One lab demonstrated that mitophagy can be carried out with BNIP3, but under conditions of BNIP3 overexpression and still involved Parkin^49^.

Based on the gene expression changes in DOX-exposed cardiac fibroblasts quantified in this study, it is possible that these cells are undergoing a senescent transformation. A recent study described a “survival” phenotype that cardiac fibroblasts undergo in response to oxidative stress^50^. Regardless of how the cells are characterized, they will likely be unable to respond adequately to future cellular and cardiac stresses. With a decreased ability to proliferate and migrate, as noted in our data, the heart will be dependent on non-cardiac sources of fibroblasts if injury arises. Furthermore, in a heart that is not fully mature, i.e. a pediatric patient, the DOX-induced cardiac fibroblast phenotype may alter the maturation of the heart. Our data suggest that early changes in the gene expression indicate ECM remodeling, with an increase in remodeling genes and a decrease in structural genes. While CFs are directing cardiac remodeling during maturation, they also participate in signaling myocyte hypertrophy. The effects of increased inflammatory cytokines in the microenvironment on long-term cardiac function are not fully known but are associated with DCM. Disruptions to this process could have long-reaching implications on cardiac function.

Out of 160 genes measured, three genes that were upregulated were cadherin-2 (Cdh2), C-X-C Motif Chemokine Ligand 10 (Cxcl10), and CxCl11. Interestingly, Cxcl10 and Cxcl11 protein are both upregulated in patients with heart disease^51; 52^ and some studies indicate this is true for DOX cardiotoxicity as well^53; 54^. Cdh2 was demonstrated to increase in dilated cardiomyopathies^55; 56^. Its role as a possible biomarker for DOX cardiotoxicity has not been investigated, though other studies have shown its gene expression is upregulated in DOX-exposed MCF-7 cells^57^. An early response to DOX-induced damage in the cardiac fibroblasts of patients may be the release of Cxcl10, Cxcl11, or Cdh2. These molecules should be further investigated as clinical prognosticators of DOX-induced cardiac disease.

Understanding the effect of DOX on a cell type vital to cardiac maturation and function may uncover more avenues of possible targets. Because cardiac fibroblasts have historically not been a major focus in research on the detrimental effects of DOX on the heart, the landscape of DOX’s effect on cardiac fibroblasts offers fertile ground for discovering novel ways to better protect the myocardium during and after DOX treatment. The use of small-molecule p53 inhibitors are currently under investigation in clinical trials for cancer therapy. In tumors without p53 deletions or mutations, p53 can actually be protective^58^. By inducing cell cycle arrest, p53 will prevent the tumor cell from proliferating or induce apoptosis. In other cases, gain-of-function mutant p53 forms can interfere with autophagy and promote anti-apoptotic pathways^59^. In these instances, the mutant p53 actually promotes tumor proliferation. Theoretically, by knocking down p53 expression during cancer treatment in specific tumors, you may sensitize the tumor to chemotherapy while protecting the cardiac fibroblasts, and thereby the heart, from disrupted mitophagy and dysfunction. Additionally, just as there is research into targeted delivery of chemotherapeutic agent to tumors, research could look into the targeted delivery of protective agents to the heart. In this way, patients who would not benefit from a p53-inhibitory treatment could still benefit from the possible cardioprotective effects of p53 inhibition.

The current study is limited by the use of mono-cultured cells. Cardiac fibroblasts and cardiac myocytes highly influence the behavior of the other. Future studies will include adapting the research to co-cultures of cardiac fibroblasts and myocytes to enhance the understanding of cell-cell communication and determine the effect of the dysfunctional fibroblast on myocyte health. An in-depth look at the role of mitochondrial dysfunction in the cardiac fibroblast is also necessary. Investigating retrograde nuclear signaling among other mitochondrial signaling mechanisms will help determine the exact connection between mitochondrial dysfunction and the DOX-induced phenotype. This study focused on only one mechanism of DOX damage (mitochondrial) and one role of p53. Future studies will also elaborate on other DOX-induced cellular damages and how an increase in p53 expression could affect other p53-involved pathways. A relevant example is the role of p53 in cellular senescence^60^. In an MI model, p53 upregulation was associated with cardiac fibroblast senescence and increased cytokine production. Ablation of p53 led to a decrease in senescent fibroblasts and increased collagen production, a protective mechanism in this situation. Other therapeutic options include salvaging mitophagy by replacing or bypassing Parkin. The BNIP3 pathway, thought to be more active during homoestatic mitophagy, could be stimulated to offset the Parkin sequestration.

The current study would best be applied to acute changes that may be seen during the early treatment phases of a cancer patient. Mitochondrial QC mechanisms occur on a very short timescale, as evidenced by activities such as exercise. Phenotypic effects of aberrant mtQC shown in this paper would then most likely be evident shortly thereafter.

These studies have demonstrated that DOX does alter cardiac fibroblast function and this has broad implications on the cardiac microenvironment and therefore cardiac function. To date, no study has investigated the effect of DOX exposure on cardiac fibroblasts in depth. The novel findings indicate an important role for fibroblasts to play in DOX cardiotoxicity that may be mediated by mitochondrial health and quality control. Increased study of the cardiac fibroblast will increase our understanding of the disease progression and create new avenues for therapeutic targets. If phenotypic changes in the cardiac fibroblast are an early indicator of cardiac toxicity, it is also possible that associated biomarkers may be developed to screen cancer drugs in development for cardiotoxic effects. If further study of the cardiac fibroblast can help protect the heart during DOX exposure, we will be able to improve the quality of life for the ever-growing population of childhood cancer survivors.

